# Hypermutator *Pseudomonas aeruginosa* exploits multiple genetic pathways to develop multidrug resistance during long-term infections in the airways of cystic fibrosis patients

**DOI:** 10.1101/809319

**Authors:** C.A. Colque, A.G. Albarracín Orio, S. Feliziani, R.L. Marvig, A.R. Tobares, H.K. Johansen, S. Molin, A.M. Smania

## Abstract

*Pseudomonas aeruginosa* exploits intrinsic and acquired resistance mechanisms to resist almost every antibiotic used in chemotherapy. Antimicrobial resistance in *P. aeruginosa* isolated from cystic fibrosis (CF) patients is further enhanced by the occurrence of hypermutator strains, a hallmark of chronic CF infections. However, the within-patient genetic diversity of *P. aeruginosa* populations related to antibiotic resistance remains unexplored. Here, we show the evolution of the mutational resistome profile of a *P. aeruginosa* hypermutator lineage by performing longitudinal and transversal analyses of isolates collected from a CF patient throughout 20 years of chronic infection. Our results show the accumulation of thousands of mutations with an overall evolutionary history characterized by purifying selection. However, mutations in antibiotic resistance genes appear to be positively selected, driven by antibiotic treatment. Antibiotic resistance increased as infection progressed towards the establishment of a population constituted by genotypically diversified coexisting sub-lineages, all of which converged to multi-drug resistance. These sub-lineages emerged by parallel evolution through distinct evolutionary pathways, which affected genes of the same functional categories. Interestingly, *ampC* and *fstI*, encoding the β-lactamase and penicillin-binding protein 3, respectively, were found among the most frequently mutated genes. In fact, both genes were targeted by multiple independent mutational events, which led to a wide diversity of coexisting alleles underlying β-lactam resistance. Our findings indicate that hypermutators, apart from boosting antibiotic resistance evolution by simultaneously targeting several genes, favor the emergence of adaptive innovative alleles by clustering beneficial/compensatory mutations in the same gene, hence expanding *P. aeruginosa* strategies for persistence.

**IMPORTANCE:** By increasing mutation rates, hypermutators boost antibiotic resistance evolution by enabling bacterial pathogens to fully exploit their genetic potential and achieve resistance mechanisms for almost every known antimicrobial agent. Here, we show how co-existing clones from a *P. aeruginosa* hypermutator lineage that evolved during 20 years of chronic infection and antibiotic chemotherapy, converged to multidrug resistance by targeting genes from alternative genetic pathways that are part of the broad *P. aeruginosa* resistome. Within this complex assembly of combinatorial genetic changes, in some specific cases, multiple mutations are needed in the same gene to reach a fine tuned resistance phenotype. Hypermutability enables this genetic edition towards higher resistance profiles by recurrently targeting these genes, thus promoting new epistatic relationships and the emergence of innovative resistance-conferring alleles. Our findings help to understand this link between hypermutability and antibiotic resistance, a key challenge for the design of new therapeutic strategies.

## INTRODUCTION

Antibiotic resistance has emerged as a global health concern with serious economic, social and political implications. Accordingly, it is becoming widely accepted that we are close to a post-antibiotic era due to the increasing occurrence of multidrug-resistant pathogens and the failure to compensate this phenomenon with drug discovery (1).

Among high-risk pathogens, *Pseudomonas aeruginosa* is one of the leading causes of nosocomial infections and the third most common bacterium isolated from infections acquired in intensive care units (2). Likewise, *P. aeruginosa* chronically infects the airways of cystic fibrosis (CF) patients and constitutes their main cause of morbidity and mortality (3).

The effective intrinsic and acquired resistance mechanisms of *P. aeruginosa* to different types of antibiotics (4) and the emergence of multidrug-resistant (MDR) clones (5), severely compromise the treatment of these infections. Notably, several resistance genes, including different classes of carbapenemases, have spread among an increasing number of *P. aeruginosa* clones through horizontal gene transfer. In many cases, this makes colistin, and to some extent amikacin, the only available drugs to treat MDR *P. aeruginosa* infections (2).

Intrinsic mechanisms of resistance involve mutations in chromosomal genes leading to the inactivation of the carbapenem porin OprD, the overexpression of AmpC and the upregulation of efflux pumps (4, 6, 7). Importantly, the concomitant accumulation of these mutations can lead to the emergence of MDR strains, which constitute a major concern in clinical setting (5).

Frequently, the acquisition of these adaptive mutations is enhanced by increments in mutation rates like that observed in hypermutator strains of *P. aeruginosa*. Thus, it has been reported that 36-54% of chronically infected CF patients are infected with hypermutator strains of *P. aeruginosa*, which are deficient in the DNA mismatch repair (MMR) system (8-13). The hypermutator phenotype has been correlated with increased development of antibiotic resistance (8, 12, 14-16), acquisition of chronic infection adaptive variants (16-19), as well as metabolic adaptive transformations (20).

Recent advances in whole-genome sequencing (WGS) techniques have provided insights into the evolutionary trajectories of adaptation of *P. aeruginosa* to the CF environment, particularly with regard to patho-adaptive mutations, such as those associated with antibiotic resistance (11, 21-26). In this sense, “the mutational resistome” was recently defined as the set of mutations involved in modulation of antibiotic resistance levels in absence of horizontal gene transfer (27, 28).

In a previous investigation, we studied the evolutionary trajectories of *P. aeruginosa* hypermutator lineages in long-term CF chronic infection (24). Comparative WGS analyses showed extensive within-patient genomic diversification, with populations composed of different sub-lineages that had coexisted for many years since the initial colonization of the patient. Importantly, certain genes were particularly enriched for mutations and underwent convergent evolution across the sub-lineages, suggesting that they are involved in the optimization process of the *P. aeruginosa* pathogenic fitness. Here, we characterize the mutational resistome and the antibiotic susceptibility profile of a hypermutator lineage sampled throughout a period of 20 years of evolution from the airways of a CF patient. To gain a comprehensive picture of the evolution of antibiotic resistance, we performed a longitudinal analysis by exploring WGS data of three sequentially isolated clones and a transversal study on a collection of 11 isolates obtained from a single sputum sample, which provided a snapshot of the genetic diversity at population level.

## RESULTS

### Emergence of multidrug-resistant isolates in the CFD collection

In our previous study, we sequenced whole genomes of 14 isolates belonging to the same clonal lineage of *P. aeruginosa*, spanning 20 years of a patient’s infection history (referred to as patient CFD) (24). This collection included a normo-mutator isolate obtained in 1991, which we used as the ancestral reference, two hypermutator isolates one from 1995 and the other from 2002, and 11 isolates obtained from the same sputum sample in 2011.

All the hypermutator isolates harbor the same *mutS* mutation, which inactivates the MMR system (24). As shown in Fig. 1, patient CFD received prolonged treatment with a large and varied set of antibiotics during the course of chronic infection between 1986 and 2012. Treatment included five classes of antibiotics: β-lactams, aminoglycosides, quinolones, polymyxins and macrolides. To investigate the impact of antibiotic treatment on the resistance profiles of the CFD isolates, susceptibility to antibiotics representing all these major classes was tested by the agar diffusion method according to the CLSI guidelines. As observed in Fig. 2, all except for the 1991 isolate showed multidrug-resistance, meaning a reduced susceptibility to two or more classes of antibiotics. Starting from the general susceptible phenotype of the 1991 isolate, the 1995 exhibited resistance to β-lactams, whereas the 2002 isolate, in addition to β-lactams, gained resistance to ciprofloxacin, tobramycin, and colistin. Importantly, the collection of 2011 isolates showed the highest levels of resistance to ciprofloxacin, tobramycin, azithromycin, colistin and particularly to β-lactams such as cephalosporins and the monobactam aztreonam. Interestingly, in contrast to the 1995 isolate, the 2002 and all the 2011 isolates showed susceptibility to piperacillin-tazobactam, resistance to which seems to have been lost after the acquisition of resistance to tobramycin, thus suggesting collateral sensitivity to penicillin-type-β-lactams as previously described by Barbosa et al. (29). On the other hand, even though colistin was used from 2004, all isolates showed MIC values ranging from 8 to 32 µg/ml, which are relatively high compared to other data sets from clinical isolates (27, 28, 30, 31) (Fig. 2). Finally, although *P. aeruginosa* has no clinical breakpoints established for azithromycin, the evolved isolates showed relatively higher MICs than the 1991 and 1995 isolates.

**Figure 1.**
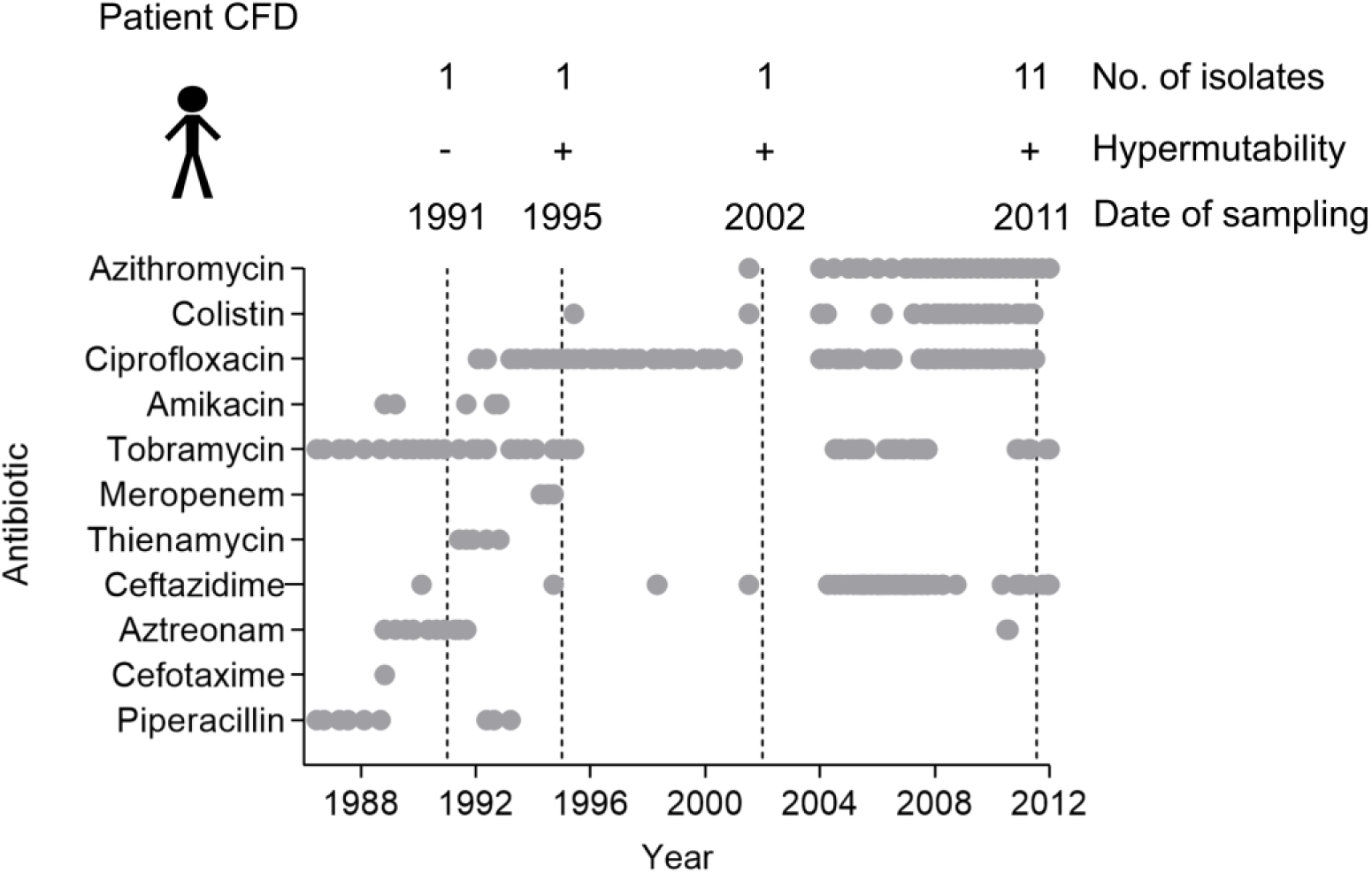
Overview of isolate sampling time points and antibiotic treatment. *P. aeruginosa* isolates were collected from patient CFD between 1991 and 2011. +/-symbols indicate hypermutability state of *P. aeruginosa* strains. Antibiotics used in chemotherapy through the 20 years study are listed in the Y axis. Grey circles indicate the start and end of an antibiotic dose.

**Figure 2.**
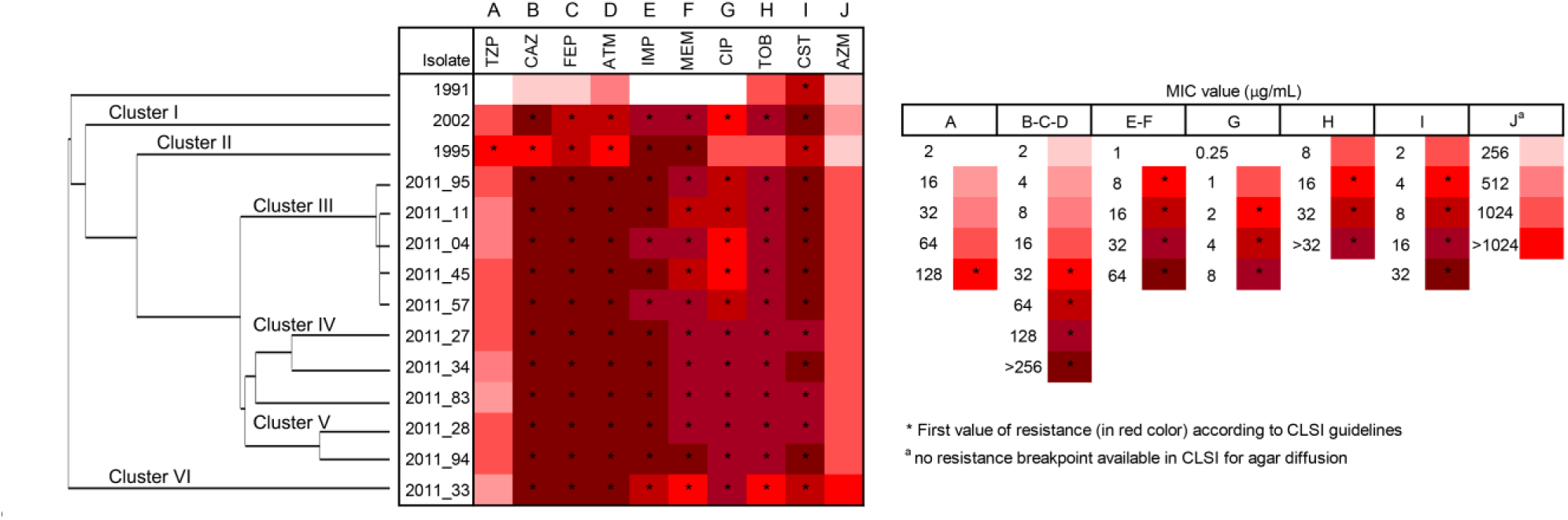
Antibiotic resistance profiles of *P. aeruginosa* isolates from the CFD lineage. Each column represents the Minimal Inhibitory Concentration (MIC) values of the different antibiotic tested: piperacillin-tazobactam (TZP); ceftazidime (CAZ); cefepime (FEP);; aztreonam (ATM); imipenem (IMI); meropenem (MEM); ciprofloxacin (CIP); tobramycin (TOB); colistin (CST) and azithromycin (AZM). Red intensity indicates MIC levels for each antibiotic. Asterisks (*) indicate resistance according to CLSI. Left tree represents the genetic clustering of isolates (rows) based on the result of maximum-parsimony analysis. The phylogenetic tree on the left was constructed based on the accumulation of new SNPs relative to ancestor 1991 (24).

### Mutations for antibiotic resistance are positively selected during evolution

In order to investigate the molecular bases of the antimicrobial resistance observed in CFD isolates, we explored the acquisition of mutations in a set of 168 chromosomal genes, here defined as the resistome, which have been described to be involved in *P. aeruginosa* antibiotic resistance mechanisms (27, 28, 32). Thus, using the 1991 genome as reference, we analyzed the distribution of a total of 5710 SNPs and 1078 indels accumulated in a period of 20 years of infection, which we have previously detected in the collection of isolates by WGS analysis (24). Furthermore, we also analyzed 39 SNPs (26 synonymous, 13 non-synonymous) detected in the genome sequence of the 1991 isolate when compared to the PAO1 genome. Sequence variations found within the resistome are documented in Supplementary Table 1 (Table S1). Interestingly, 93 (55%) of the 168 investigated genes, showed non-synonymous SNPs and/or indels mutations of 1-3 bp in at least one of the isolates. On the other hand, 11 (6%) genes showed only synonymous mutations, and 64 (38%) showed no mutations (Fig. S1). Furthermore, 86 out of the 87 genes harboring non-synonymous mutations (99%), were targeted with missense mutations, whereas a single gene (1%) showed a nonsense mutation (Fig. S1 and Table S1). By analyzing the ratio between non-synonymous and synonymous mutations (dN/dS ratio) within the resistome in each CFD isolate (Table S2), we observed that in most isolates the signature of selection was higher than 1 and higher than the ratio obtained from SNPs affecting all the other genes (dN/dS=0.78). This indicates that these mutations were positively selected during chronic infection, and suggests that hypermutability may be linked to them as a key factor contributing to antibiotic resistance in CF. Mutational resistome analysis was further focused on those genes that were targeted with non-synonymous and/or frameshift mutations (Fig. 3), showing that accumulation of these mutations correlated with increased antibiotic resistance. Moreover, no mutations were detected in the ancestral 1991 isolate respect to the reference strain PAO1, in agreement with its general antibiotic susceptibility (Fig. 2). In some genes known to be involved in antibiotic resistance, single mutations were identified, such as D87G in GyrA and S278P in OprD (33, 34); in others, the accumulation of 3 to 5 different mutational events suggests that they have evolved under strong selective pressure. Such examples are *amgS, mexX, fusA2* (involved in aminoglycoside resistance), *mexF, oprN, poxB, mexI* (involved in β-lactams resistance), *polB, mexD, parE* (involved in quinolone resistance), *spuF* (polyamines) and *mexK* (coding for a novel efflux system MexJK) (35). Remarkably, *mexY, fusA1* (aminoglycoside resistance), *ampC* and *ftsI* (β-lactams resistance) accumulated more than 6 different mutations (Fig. 3), providing strong evidence for parallel evolution.

**Figure 3.**
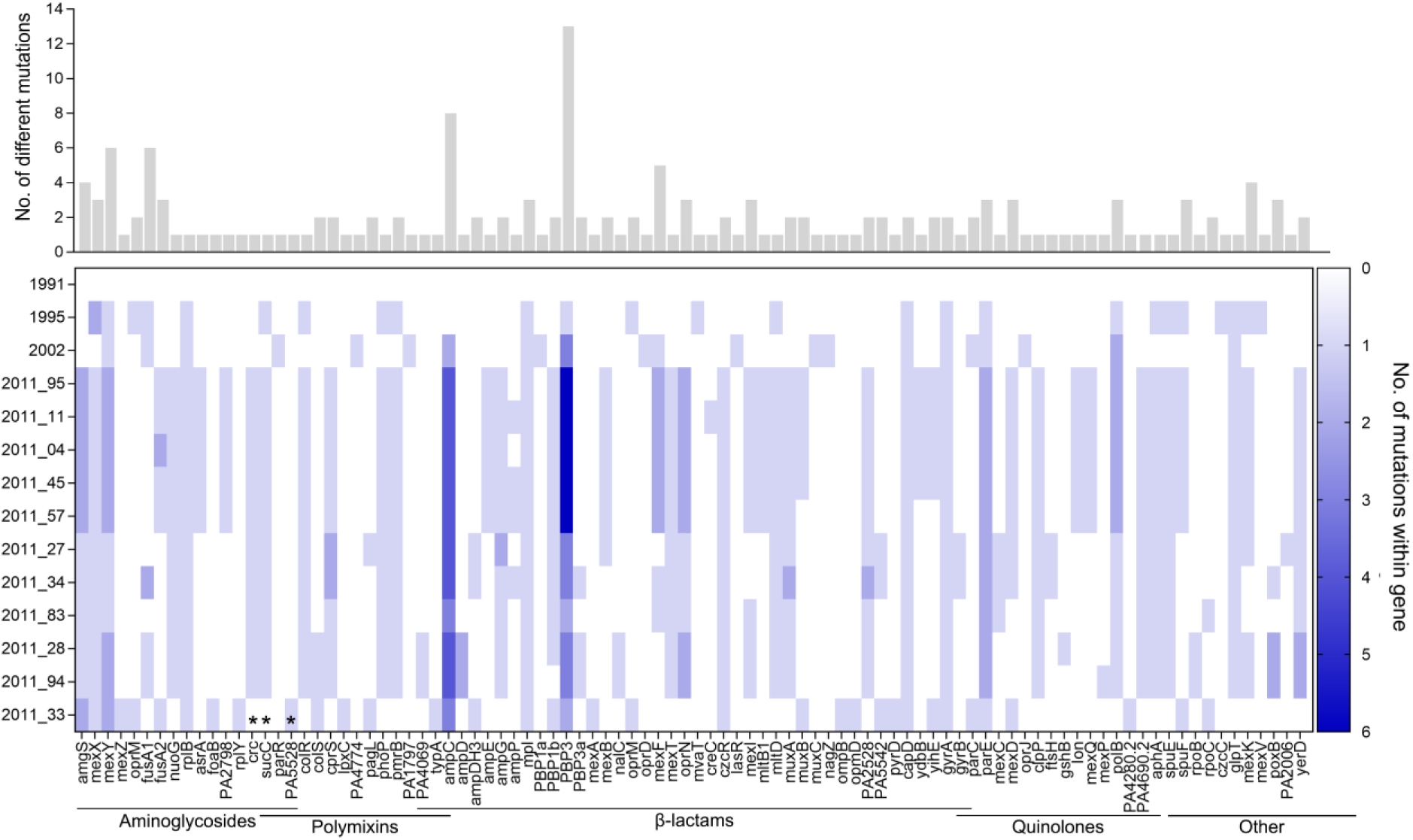
Resistome of the CFD isolate collection. Mutations potentially affecting protein function in 93 out of the 168 antibiotic resistance genes were analyzed. Genes and variants were grouped by antibiotic class. Upper panel: number of independent mutations found within each specific gene along the CFD lineage. Lower panel: heatmap of the number of mutations accumulated *per* gene in each genome. Isolates were grouped based on the genetic clustering defined in Fig. 2. *Asterisks indicate genes which are also involved in conferring resistance to other antibiotic classes; *crc* (β-lactams), *sucC* and *PA5528* (quinolones).

### The β-lactam resistome

As shown in Fig. 1, patient CFD received prolonged antibiotic courses with Ceftazidime as well as shorter courses of varying durations with different types of β-lactams, including other cephalosporins (Cefotaxime), penicillins (Piperacillin + tazobactam), monobactams (Aztreonam) and carbapenems (Thienamycin and Meropenem). As expected, resistance increased from the 1995 isolate to later isolates, reaching the highest resistance levels to cephalosporins, aztreonam and carbapenems in the 2011 isolates (Fig. 2). A total of 70 genes have been reported to be involved in the β-lactam resistome (27, 28), including: regulation of peptidoglycan-recycling genes (responsible for AmpC overproduction), genes encoding penicillin-binding proteins (PBPs, targets of β-lactam antibiotics), or encoding regulators of efflux pumps such as mexAB-oprM (involved in β-lactam resistance) and mexEF-OprN (involved in carbapenem resistance) and, the *oprD* gene (involved in resistance to imipenem and susceptibility to meropenem). We found that 42 of these 70 genes (60%) showed non-synonymous and/or frameshift mutations in at least one isolate of the CFD collection (Fig. 3). Of these 42 genes, 36 showed accumulations of 1 to 2 mutations, most of them being unique to each specific cluster. It has been described that the emergence of resistance to penicillins and cephalosporins is mainly due to overproduction of the β-lactamase AmpC (36). However, the most frequent drivers of AmpC overproduction described in *P. aeruginosa* clinical strains, namely *ampD, ampR* and *dacB* (37-39), were not mutated among CFD isolates. Instead, all but the 1991 isolate showed a frameshift mutation in *mpl*, which encodes a UDP-N-acetylmuramate:l-alanyl-γ-d-glutamyl-meso-diaminopimelate ligase, indicating that *ampC* could be overexpressed via this alternative negative regulator (40). Western blot analyses showed that all 2011 isolates showed an increased expression of AmpC compared to the 1991, 1995 and 2002 isolates, suggesting that alternative pathways may be responsible for AmpC overproduction in these isolates (Fig. S2). In addition, *ampC* was among the most mutated genes in the CFD collection together with the *ftsI* gene, showing 8 and 13 distinct missense mutations, respectively (Fig. 3). In fact, all CFD isolates, except 1991 and 1995, showed accumulation of mutations within *ampC*, with isolates from 2011 carrying up to four different mutations, combined in different *ampC* alleles. This strongly correlates with the increase in the MICs of cephalosporins and with aztreonam resistance in the evolved CFD isolates (Fig. 2), indicating that they were under high selective pressure during the CFD chronic infection process. Interestingly, the presence of mutations such as P154L, G216S and V213A, has been reported to be involved in β-lactam resistance (41). Likewise, all isolates except for 1991 showed mutations in *ftsI*, compared with isolates from 2011 showing up to six mutations combined in a single allele. Some of these mutations (Y367C, H394R, N427S, Q458R, Q475R, R504L, V523A, V523M and F533L) are located in the transpeptidase β-lactam binding site of the protein, and the latter mutation has been shown to play a key role in β-lactam recognition (42). Importantly, these mutations have been documented to emerge among *P. aeruginosa* CF collections (28, 43, 44) and upon aztreonam exposure *in vitro* (45), whereas other mutations are described for the first time in this work (Table S1). The rest of the PBP encoding genes showed few mutations among the CFD collection.

Although patient CFD received only short courses with carbapenems, we observed the emergence of high levels of resistance to carbapenems in all isolates except for the ancestral 1991 isolate. Previous reports have shown that loss of function mutations in the outer membrane protein OprD and/or overexpression of efflux pumps MexAB-OprM (meropenem resistance) and MexEF-OprN (imipenem and meropenem resistance) constitute main mechanisms to develop carbapenem resistance. However, only the 2002 isolate (Cluster I, defined according the phylogenetic tree showed in Figure 2) showed a missense mutation (S278P) within the *oprD* gene, which has been previously described to be involved in carbapenem resistance (34). Expression of the MexAB-oprM system is controlled by the regulatory genes *mexR, nalC* and *nalD* (46, 47), and a missense mutation in *nalC* (M151T) was identified in the two isolates from Cluster V. Mutations F533L and R504 in PBP3 have been found to occur upon meropenem exposure during *in vitro* evolution studies and among CF patients treated with this drug (44, 48). Thus, high levels of carbapenem resistance may be associated with the presence of these *ftsI* mutations. Importantly, using ResFinder tool on WGS data from CFD isolates, we did not find genes coding for any class of β Metallo-beta-lactamases (MBLs) involved in carbapenem resistance, which are normally acquired through horizontal gene transfer (49). These results suggest that various different mutational mechanisms may be involved in carbapenem resistance in different coexisting CFD isolates, giving rise to distinct genetic pathways for the evolution of resistance to β-lactams.

### The aminoglycoside resistome

As shown in Fig. 1, patient CFD received extensive treatment courses of tobramycin. MIC determinations showed that all isolates, except for 1991 and 1995, became resistant to tobramycin. The main origin of high-level resistance to aminoglycosides is the overexpression of MexXY-OprM efflux system (50), which is primarily caused by *mexZ* mutations (10, 21, 51). In addition, mutations in the *amgRS* and *parRS* two-component systems genes have also been involved in the regulation of MexXY expression (52). No *mexZ* mutations, however, were observed in the CFD collection of isolates, with the sole exception of isolate 2011_33 (Cluster VI), which showed a V29A mutation located within the DNA binding domain of the protein (Table S1) (53) and predicted to be deleterious (−3.351 PROVEAN v1.1.3). Instead, we found four different mutations in gene *amgS*, encoding the histidine kinase sensor of the membrane stress-response two-component system, six mutations in *mexY*, encoding a component of the MexXY efflux pump, and six mutations in *fusA1*, which codes for the elongation factor G.

MexY mutations have been frequently observed among drug-resistance isolates and CF epidemic clones (28, 31). Some mutations affect the general pump operation and impair the MexY-dependent aminoglycoside resistance, whereas other mutations, located in domains associated with aminoglycoside recognition and export, may improve drug accommodation and consequently increase resistance (54). Furthermore, it was observed that the MexY mutation F1018L is able to increase pump-promoted resistance to aminoglycosides, cefepime, and fluoroquinolones (55). Importantly, here we describe for the first time the six *mexY* missense mutations. In this sense, their impact in MexXY pump function and aminoglycoside resistance remains unclear and deserves further investigation.

Mutations in the *amgS* gene have been shown to be involved in intrinsic aminoglycoside resistance in *P. aeruginosa* (56). Although none of the four mutations in *amgS* found here have been previously reported (Table S1), all isolates from 2011 except 2011_33 showed an A203V mutation located within the linker HAMP domain. Interestingly, it has been reported that mutations in the linker domain of EnvZ, the closest *E. coli* homolog of AmgS, often cause activation of the kinase sensor (57, 58). Moreover, we found the P116L mutation, predicted to be deleterious (−2.539 PROVEAN v1.1.3). This mutation is located in the sensor domain of AmgS, where mutations involved in aminoglycoside resistance have been previously described (56).

FusA1 mutations have been recently linked to the emergence of aminoglycoside resistance *in vitro* (59-61) as well as in clinical CF strains (28, 62-64). In fact, aminoglycoside resistance seems to be an indirect consequence of the alteration of elongation factor G (60). Isolate 2011_34 harbored two substitutions, V93A and D588G, located in domains G and IV of the protein, respectively, which have been reported to be gain-of-function mutations (28). Indeed, the V93A mutation was found to increase resistance to several aminoglycosides such as tobramycin, amikacin, and gentamycin (60). In several CFD isolates we identified four novel mutations in the *fusA1* gene across the different domains of the protein sequence: domain II (V338A), domain III (A481V), domain IV (A595V) and domain V (Y683C). Isolate 1995, harboring the Y683C mutation, showed susceptibility to tobramycin (Fig. 2), suggesting that this mutation is not involved in aminoglycoside resistance. Moreover, Bolard et al. (60) recently reported that higher MICs are associated with mutations in domains II, IV and V, but not in domains G and III. Therefore, only mutations V338A (isolate 2002) and A595V (isolates 2011 from cluster III; Fig. 2) are expected to contribute to aminoglycoside resistance, although the effect of both substitutions needs to be characterized in future works. In conclusion, high-level of aminoglycoside resistance in the CFD population seems to have been acquired mostly by different mutations in the *amgS* and/or *fusA1* genes.

### The fluoroquinolone resistome

Patient CFD received two prolonged periods of treatment with ciprofloxacin, from 1992 to 2002 and from 2004 to 2012 (Fig. 1). MICs of ciprofloxacin revealed that most of the isolates exhibited high resistance levels to this antibiotic, whereas the 1991 and 1995 isolates showed susceptibility and intermediate resistance, respectively (Fig. 2). High resistance to ciprofloxacin usually involves one or several mutations in quinolone resistance determining (QRD) regions of the GyrAB subunits of topoisomerase II (gyrase), and the ParCE subunits of topoisomerase IV (28). Indeed, all CFD isolates except for the 1991, harbored the same D87G mutation in GyrA. In addition, isolate 2011_33 also carried a T83I mutation in this gyrase subunit. Importantly, both mutations are known to be involved in quinolone resistance (28, 31, 33). Furthermore, two 2011 isolates from Cluster IV harbored an S618L substitution in GyrB. On the other hand, all isolates except for the 1991, accumulated mutations in the topoisomerase IV subunits ParC (P308L, T705A) and ParE (V199M, D462G, S492F), none of which have been previously described. Whether these mutations clustered in the chromosomally encoded topoisomerases II and IV were involved in quinolone resistance or were randomly fixed by genetic drift upon the high mutation supplies provided by hypermutability, remains to be elucidated. Nevertheless, the fact that many different mutations arose after fluoroquinolone treatments supports the previous observation that mutations involved in fluoroquinolone resistance can be highly variable (28). Importantly, with the use of the ResFinder tool we found the acquisition of a novel plasmid-encoded ciprofloxacin-modifying gene encoding the enzyme CrpP (65), which may explain the high-resistance profile observed in the two intermediate isolates from 1995 and 2002, and all isolates from 2011.

Finally, we noticed that no mutations were observed in the negative regulator *nfxB* among the CFD isolates, which is commonly reported to achieve resistance to ciprofloxacin in a CF context due to the deregulation and concomitant overexpression of the efflux pump MexCD-OprJ (66). Furthermore, although all Cluster IV isolates from 2011 harbored a nonsense mutation in the transporter MexD (W1023STOP), which inactivates the efflux pump, it has been described that this mutation has no effect on the MIC of ciprofloxacin (67).

### The polymyxin resistome

Patient CFD received intensive treatment with colistin from 2004 to 2011 (Fig. 1). According to CLSI, antibiotic susceptibility profiling revealed that every CFD isolate was resistant to colistin (Fig. 2). However, the evolved 2011 isolates from cluster III, IV and V as well as the 2002 isolate, showed 2- to 4-fold increases in their MICs relative to 1991 and 1995 isolates (8 μg/mL) (Fig. 2). Clinical strains of *P. aeruginosa* sometimes show resistance to polymyxins due to mutations in different two-component systems, such as PhoPQ, PmrAB, ParRS, CprRS and ColRS (68-72). Additionally, mutations causing derepression of the lipopolysaccharide (LPS) modifying (*arn*) operon, encoding the proteins necessary for the aminoarabinosylation of the lipid A moiety of the LPS, have been identified in colistin-resistant *P. aeruginosa* strains (30, 70, 73). As shown in Fig. 3, the different CFD isolates accumulated unique mutations in genes *phoP, pmrB, parR, colR* and *colS* genes, which may affect each of the mentioned two-component systems. In fact, the 2002 isolate harbored a A45T mutation in ParR located in the receiver domain and close to the conserved phosphorylation residue D57, which was previously shown to be involved in colistin resistance (74). On the other hand, considering that mutations V30A in PhoP, and D138N in ColR were present in the more susceptible 1995 isolate, the increased resistance observed in the 2011 evolved isolates from clusters III, IV and V could be explained by the presence of mutations in PmrB (T132A), CprS (G396S), and/or ColS (T138A). These novel mutations are the first to suggest their contribution to polymyxin resistance and therefore need to be further explored.

### Other antibiotics

From the beginning of 2004 to 2012, patient CFD received systematic long-term treatments with azithromycin combined with other antipseudomonal agents. Although macrolide resistance is frequent among CF isolates, only two reports describe the emergence of macrolide resistance *in vivo* (32, 75). As shown in Fig. 2, MICs of azithromycin for the later isolates within the 2011 collection showed a 4-fold increase or more, relative to the ancestral isolate from 1991. Consistent with this, all 2011 isolates carried mutations in gene PA4280.2, which encodes the 23S ribosomal subunit. In fact, isolate 2011_33 carried an A2044G substitution, whereas the remaining isolates from 2011 carried a C2597T mutation, both located in the secondary structure of domain V of the ribosomal RNA gene and previously reported to confer macrolide resistance (32, 75). Thus, macrolide resistance in coexisting CFD 2011 isolates was acquired by distinct mutations in the same gene. This provides additional evidence for parallel molecular evolution at population level, with antibiotic chemotherapy as the key selection force during long-term CF chronic infections.

## DISCUSSION

The high prevalence of hypermutator clones in CF chronic infections is a matter of great relevance because their link to antibiotic resistance hampers infection management (8-10, 14, 24, 76). In this study, we explored the evolution of the mutational antibiotic resistome of a *P. aeruginosa* hypermutator lineage by combining a longitudinal and a transversal analysis that covered 20 years of CF chronic infection. Antibiotic resistance increased as infection progressed towards the establishment of a population consisting of genotypically diversified coexisting sub-lineages, all of which converged to multi-drug resistance. Particularly, while mutations observed in *amgS* are most likely altering the MexXY pump regulation, mutations affecting other multi-drug efflux pump regulators were only rarely observed among CFD isolates. Instead, multidrug resistance emerged through the combination of multiple resistance mutations in several independent loci. Hypermutators can be indirectly selected for and fixed by their genomic association with fitness-improving alleles (77-80). Particularly, under selective conditions imposed by long-term antibiotic therapy in the CF airways, *de novo* beneficial mutations can be expected to accumulate over time. Early mutational events occurring during the course of long-term infection are expected to have a strong impact on the resistance phenotype and consequently on fitness. Later mutations, many of them compensatory, may lead to fine tuning of the activity/stability of the resistance related proteins, in which epistatic interactions may play important roles for the trajectories of resistance development (81-84). *P. aeruginosa* carries many different genes, which upon functional mutations provide a resistance phenotype (27, 28). Identification of these genes and the associated polymorphisms involved in resistance document how many of them converge through distinct genetic pathways to the same or similar resistance profiles (26, 85, 86).

Hypermutability increases the likelihood of reaching the most appropriate combinations in adaptive terms. Considering the multiple genetic pathways in *P. aeruginosa* behind different resistance mechanisms, our observations show how hypermutability increases the probability of exploiting these distinct pathways, which eventually converge towards antibiotic multi-resistance in the course of long-term chronic infections.

We show how aggressive and persistent chemotherapy targeting a hypermutator population resulted in repeated but independent mutagenic events in resistance associated genes, providing clear evidence of parallel evolution in clones of the CFD population. This was for example the case with *ampC* and *fstI*, which for the CFD lineage constituted hot spots for the accumulation of mutations involved in β-lactam antibiotic resistance (39, 41, 48, 49). Some of these mutations have been previously described, whereas others are reported here for the first time. Most interestingly, novel alleles were observed, each harboring a combination of 2 to 6 mutations.

Overproduction of the β-lactamase AmpC is considered to be the main cause of resistance to first- and second-generation cephalosporins as well as aminopenicillins in *P. aeruginosa* clinical strains (36). However, *P. aeruginosa* is also able to adapt to new and more effective β-lactams (87), through a variety of mutations affecting the AmpC β-lactamase (41, 48, 88). Here, we document the confluence of both strategies: variants overproducing AmpC, in which the combination of distinct mutations may contribute to even higher levels of resistance and/or substrate spectrum extension. Furthermore, the accumulation of several different mutations in the penicillin-binding protein PBP3 may be a complementary and/or additional pathway. PBP3 relevance in β-lactam resistance, including the new generation cephalosporins and carbapenems, has been very recently confirmed (27, 28, 31, 43, 48). The high number of different mutations clustered in both *ampC* and *fstI* genes combines into innovative resistance-conferring alleles, which demonstrate how drug-resistance mutations can become highly beneficial when combined with compensatory mutations, and thus document the extraordinary ability of *P. aeruginosa* to develop antibiotic resistance. In this context, the emergence of such innovative alleles may be distinctly favored by hypermutator phenotypes, then limiting our available therapeutic arsenal.

How can we understand co-existence of many different genetic variants showing the same resistance profile in patient airways, such as it has been documented here? One answer is based on the balance between clonal interference and multiple mutations (89, 90). We thus argue that hypermutability increases the rate of antibiotic resistance evolution by increasing piggy-backing of multiple resistance mutations, causing maintenance of a diversified population where adaptive variation is sustained by a dynamic equilibrium between mutation and selection. Moreover, in long-term evolutionary scenarios such as chronic infections, the selective forces imposed by antibiotics along with high mutation rates from hypermutability, may shape genetically diverse populations able to respond successfully to antibiotic treatments ensuring persistence of the bacterium.

Our results provide new evidence concerning the way in which hypermutators can expedite the evolution of multidrug-resistance by increasing the probability of acquiring adaptive mutations to support long-term survival of *P. aeruginosa* in the airways of CF patients.

## MATERIALS AND METHODS

### *P. aeruginosa* CFD collection

Clinical *P. aeruginosa* isolates were obtained from sputum samples from a CF patient attending the Copenhagen CF Centre at Rigshospitalet (Copenhagen, Denmark) (patient CFD). In a previous study (24), we sequenced the genomes of 14 isolates from this patient covering ∼20 years of the patient lifespan (European Nucleotide Archive, ENA/SRA ERP002379). The CFD collection included: One normo-mutator isolate obtained in 1991 (CFD_1991) five years after the onset of chronic *P. aeruginosa* infection in 1986; two sequential mismatch repair (MRS) deficient mutators from 1995 (CFD_1995) and 2002 (CFD_2002) that harbored the same ΔCG mutation in *mutS* at position 1551; and 11 *P. aeruginosa* isolates obtained from a single sputum sample in 2011 (CFD_2011), all harboring the ΔCG *mutS* mutation at 1551 and belonging to the same hypermutator lineage.

### Profiling of antibiotic resistance genes

In order to correlate the documented resistance genotypes with the observed resistance phenotypes, single-nucleotide polymorphisms (SNPs) and indels (1- to 10-bp insertion/deletion mutations) for each isolate obtained from the previous study (24) were filtered based on an exhaustive literature review (27, 28). We also added PA0668.4, PA4280.2, PA4690.2 and PA5369.2 genes to the list, affecting macrolide resistance (32). Thus, we obtained a set of 168 genes known to be related to antibiotic resistance in *P. aeruginosa*. Indels and premature stop codons were considered to result in the inactivation of the corresponding protein product. The contribution of the documented SNPs to the phenotype was evaluated according to the available literature and by using online software tools for prediction of the effect of nucleotide substitutions on protein function, e.g. SIFT (91), PROVEAN (92) and SNAP2 (93). In addition, the online tool ResFinder v2.1 (https://cge.cbs.dtu.dk//services/ResFinder/) (94) was used to identify possible horizontally acquired antimicrobial resistance genes.

### Susceptibility testing

Cs determination was performed by using the broth dilution method, according to Clinical and Laboratory Standards Institute (CLSI) guidelines and breakpoints (95). Ten antimicrobials agents from five classes of antibiotics were tested. From the β-lactam class, shown as ≤susceptible/≥resistant breakpoints, ceftazidime (8/32µg/ml), cefepime (8/32µg/ml), piperacillin/tazobactam (16-4/128-4µg/ml), aztreonam (8/32µg/ml), imipenem (2/8 µg/ml) and meropenem (2/8 µg/ml) were used. Aminoglycosides: tobramycin (4/16µg/ml); fluoroquinolones: ciprofloxacin (0.5/2µg/ml); polymyxins: colistin (2/4µg/ml); macrolides: azithromycin (no information). *P. aeruginosa* ATCC 27853 was used as quality control strain.

### AmpC expression levels

CFD isolates were grown for 16 h on LB media and 1.5 mL of each culture was pelleted and resuspended in 20 mM Tris-HCl (pH 7.4), 0.5 M NaCl, 15% glycerol, amended with 0.2 mg/ml lysozyme, 1 mM 8 phenylmethylsulfonyl fluoride and 1 mM benzamidine, and incubating for 1 h on ice. After four sonication (2 min) and freeze/unfreeze cycles, intact cells were removed by centrifugation at 9000 g for 20 min and the extracts were stored at −20°C. 25μg of total proteins were separated through sodium dodecyl sulfate (SDS)-polyacrylamide gel electrophoresis (PAGE) 12%, then proteins were transferred to polyvinylidene fluoride (PVDF) membranes for 1.5 hours at 350 mA. The blots were blocked for one hour in 5% milk in phosphate-buffered saline (PBS) solution at room temperature. Incubation with primary antibody (rabbit anti-PDC-3 policlonal, (96), was added at 1/1,000 overnight at 4°C in 5% milk/PBS, then washings were performed with PBS/Tween 20, and the secondary antibody (IRDye 680RD anti-rabbit, LI-COR) was added at a 1:20,000 dilution for 1 hour in 5% milk/PBS. Membranes were scanned on Odyssey infrared imager instrument (LI-COR Bioscience).

## AKNOWLEDGEMENTS

We are grateful to Dr. Alejandro Moyano for the reading of the manuscript and for worthwhile discussion. This work was supported by the Agencia Nacional de Promoción Científica y Tecnológica (ANPCYT) grant PICT-2016–1545, Secretaría de Ciencia y Tecnología (SECYT-UNC) grant 33620180100413CB, and the Ministerio de Ciencia y Tecnología (MINCyT) de la Provincia de Córdoba, Argentina, PID-2018 (Res N °144).

HKJ was supported by The Novo Nordisk Foundation as a clinical research stipend (NNF12OC1015920), by Rigshospitalets Rammebevilling 2015-17 (R88-A3537), by Lundbeckfonden (R167-2013-15229), by Novo Nordisk Fonden (NNF15OC0017444), by RegionH Rammebevilling (R144-A5287) and by Independent Research Fund Denmark / Medical and Health Sciences (FSS-4183-00051). We thank Ulla Rydahl Johansen for excellent technical assistance.

## SUPPLEMENTARY LEGENDS

**Figure S1. Analysis of type of mutations found in antibiotic resistance genes in *P. aeruginosa* isolates from CFD patient**.

Pie charts indicate the observed percentage for each kind of mutation respect to the total number of mutations occurring in the 168 belonging to the *P. aeruginosa* resistome.

**Figure S2. Western blot of CFD isolates**.

Total proteins (25μg) were obtained from whole-cell lysates from each *P. aeruginosa* clinical isolates, resolved in a 12% polyacrylamide gel, and tested with a PDC-3 antibody.

**Table S1. Nonsynonymous and frameshift mutations found within 93 out of 168 antibiotic resistance genes in CFD collection**.

**Table S2. Number of genes mutated and type of mutations found in the sequenced genomes**.

## REFERENCES

1. Brown ED, Wright GD. 2016. Antibacterial drug discovery in the resistance era. Nature 529:336.

2. Sader HS, Farrell DJ, Flamm RK, Jones RN. 2014. Antimicrobial susceptibility of gram-negative organisms isolated from patients hospitalized in intensive care units in United States and European hospitals (2009–2011). Diagnostic Microbiology and Infectious Disease 78:443–448.

3. Lyczak JB, Cannon CL, Pier GB. 2002. Lung infections associated with cystic fibrosis. Clinical microbiology reviews 15:194–222.

4. Breidenstein EB, de la Fuente-Nunez C, Hancock RE. 2011. *Pseudomonas aeruginosa*: all roads lead to resistance. Trends Microbiol 19:419–26.

5. Oliver A, Mulet X, Lopez-Causape C, Juan C. 2015. The increasing threat of *Pseudomonas aeruginosa* high-risk clones. Drug Resist Updat 21-22:41–59.

6. Lister PD, Wolter DJ, Hanson ND. 2009. Antibacterial-resistant *Pseudomonas aeruginosa*: clinical impact and complex regulation of chromosomally encoded resistance mechanisms. Clin Microbiol Rev 22:582–610.

7. Poole K. 2011. *Pseudomonas Aeruginosa*: resistance to the max. Frontiers in Microbiology 2.

8. Oliver A, Canton R, Campo P, Baquero F, Blazquez J. 2000. High frequency of hypermutable *Pseudomonas aeruginosa* in cystic fibrosis lung infection. Science 288:1251–4.

9. Oliver A. 2010. Mutators in cystic fibrosis chronic lung infection: prevalence, mechanisms, and consequences for antimicrobial therapy. Int J Med Microbiol 300:563–72.

10. Feliziani S, Luján AM, Moyano AJ, Sola C, Bocco JL, Montanaro P, Canigia LF, Argaraña CE, Smania AM. 2010. Mucoidy, quorum sensing, mismatch repair and antibiotic resistance in *Pseudomonas aeruginosa* from cystic fibrosis chronic airways infections. PLOS ONE 5:e12669.

11. Marvig RL, Johansen HK, Molin S, Jelsbak L. 2013. Genome analysis of a transmissible lineage of *Pseudomonas aeruginosa* reveals pathoadaptive mutations and distinct evolutionary paths of hypermutators. PLOS Genetics 9:e1003741.

12. Ciofu O, Riis B, Pressler T, Poulsen HE, Hoiby N. 2005. Occurrence of hypermutable *Pseudomonas aeruginosa* in cystic fibrosis patients is associated with the oxidative stress caused by chronic lung inflammation. Antimicrob Agents Chemother 49:2276–82.

13. Waine DJ, Honeybourne D, Smith EG, Whitehouse JL, Dowson CG. 2008. Association between hypermutator phenotype, clinical variables, mucoid phenotype, and antimicrobial resistance in *Pseudomonas aeruginosa*. Journal of Clinical Microbiology 46:3491.

14. Macia MD, Blanquer D, Togores B, Sauleda J, Perez JL, Oliver A. 2005. Hypermutation is a key factor in development of multiple-antimicrobial resistance in *Pseudomonas aeruginosa* strains causing chronic lung infections. Antimicrob Agents Chemother 49:3382–6.

15. Montanari S, Oliver A, Salerno P, Mena A, Bertoni G, Tummler B, Cariani L, Conese M, Doring G, Bragonzi A. 2007. Biological cost of hypermutation in *Pseudomonas aeruginosa* strains from patients with cystic fibrosis. Microbiology 153:1445–54.

16. Mena A, Smith EE, Burns JL, Speert DP, Moskowitz SM, Perez JL, Oliver A. 2008. Genetic adaptation of *Pseudomonas aeruginosa* to the airways of cystic fibrosis patients is catalyzed by hypermutation. J Bacteriol 190:7910–7.

17. Moyano AJ, Lujan AM, Argarana CE, Smania AM. 2007. MutS deficiency and activity of the error-prone DNA polymerase IV are crucial for determining *mucA* as the main target for mucoid conversion in *Pseudomonas aeruginosa*. Mol Microbiol 64:547–59.

18. Lujan AM, Moyano AJ, Segura I, Argarana CE, Smania AM. 2007. Quorum-sensing-deficient (*lasR*) mutants emerge at high frequency from a *Pseudomonas aeruginosa mutS* strain. Microbiology 153:225–37.

19. Hogardt M, Hoboth C, Schmoldt S, Henke C, Bader L, Heesemann J. 2007. Stage-specific adaptation of hypermutable *Pseudomonas aeruginosa* isolates during chronic pulmonary infection in patients with cystic fibrosis. J Infect Dis 195:70–80.

20. Hoboth C, Hoffmann R, Eichner A, Henke C, Schmoldt S, Imhof A, Heesemann J, Hogardt M. 2009. Dynamics of adaptive microevolution of hypermutable *Pseudomonas aeruginosa* during chronic pulmonary infection in patients with cystic fibrosis. J Infect Dis 200:118–30.

21. Smith EE, Buckley DG, Wu Z, Saenphimmachak C, Hoffman LR, D’Argenio DA, Miller SI, Ramsey BW, Speert DP, Moskowitz SM, Burns JL, Kaul R, Olson MV. 2006. Genetic adaptation by *Pseudomonas aeruginosa* to the airways of cystic fibrosis patients. Proc Natl Acad Sci U S A 103:8487–92.

22. Yang L, Jelsbak L, Marvig RL, Damkiaer S, Workman CT, Rau MH, Hansen SK, Folkesson A, Johansen HK, Ciofu O, Hoiby N, Sommer MO, Molin S. 2011. Evolutionary dynamics of bacteria in a human host environment. Proc Natl Acad Sci U S A 108:7481–6.

23. Winstanley C, O’Brien S, Brockhurst MA. 2016. *Pseudomonas aeruginosa* evolutionary adaptation and diversification in cystic fibrosis chronic lung infections. Trends Microbiol 24:327–337.

24. Feliziani S, Marvig RL, Luján AM, Moyano AJ, Di Rienzo JA, Krogh Johansen H, Molin S, Smania AM. 2014. Coexistence and within-host evolution of diversified lineages of hypermutable *Pseudomonas aeruginosa* in long-term cystic fibrosis infections. PLOS Genetics 10:e1004651.

25. Marvig RL, Sommer LM, Molin S, Johansen HK. 2015. Convergent evolution and adaptation of *Pseudomonas aeruginosa* within patients with cystic fibrosis. Nat Genet 47:57–64.

26. Folkesson A, Jelsbak L, Yang L, Johansen HK, Ciofu O, Hoiby N, Molin S. 2012. Adaptation of *Pseudomonas aeruginosa* to the cystic fibrosis airway: an evolutionary perspective. Nat Rev Microbiol 10:841–51.

27. Cabot G, Lopez-Causape C, Ocampo-Sosa AA, Sommer LM, Dominguez MA, Zamorano L, Juan C, Tubau F, Rodriguez C, Moya B, Pena C, Martinez-Martinez L, Plesiat P, Oliver A. 2016. Deciphering the resistome of the widespread *Pseudomonas aeruginosa* sequence type 175 international high-risk clone through whole-genome sequencing. Antimicrob Agents Chemother 60:7415–7423.

28. López-Causapé C, Sommer LM, Cabot G, Rubio R, Ocampo-Sosa AA, Johansen HK, Figuerola J, Cantón R, Kidd TJ, Molin S, Oliver A. 2017. Evolution of the *Pseudomonas aeruginosa* mutational resistome in an international cystic fibrosis clone. Scientific reports 7:5555–5555.

29. Barbosa C, Trebosc V, Kemmer C, Rosenstiel P, Beardmore R, Schulenburg H, Jansen G. 2017. Alternative evolutionary paths to bacterial antibiotic resistance cause distinct collateral effects. Mol Biol Evol 34:2229–2244.

30. Fernandez L, Alvarez-Ortega C, Wiegand I, Olivares J, Kocincova D, Lam JS, Martinez JL, Hancock RE. 2013. Characterization of the polymyxin B resistome of *Pseudomonas aeruginosa*. Antimicrob Agents Chemother 57:110–9.

31. del Barrio-Tofiño E, López-Causapé C, Cabot G, Rivera A, Benito N, Segura C, Montero MM, Sorlí L, Tubau F, Gómez-Zorrilla S, Tormo N, Durá-Navarro R, Viedma E, Resino-Foz E, Fernández-Martínez M, González-Rico C, Alejo-Cancho I, Martínez JA, Labayru-Echverria C, Dueñas C, Ayestarán I, Zamorano L, Martinez-Martinez L, Horcajada JP, Oliver A. 2017. Genomics and susceptibility profiles of extensively drug-resistant *Pseudomonas aeruginosa* isolates from Spain. Antimicrobial Agents and Chemotherapy 61:e01589–17.

32. Mustafa MH, Khandekar S, Tunney MM, Elborn JS, Kahl BC, Denis O, Plesiat P, Traore H, Tulkens PM, Vanderbist F, Van Bambeke F. 2017. Acquired resistance to macrolides in *Pseudomonas aeruginosa* from cystic fibrosis patients. Eur Respir J 49.

33. Kos VN, Déraspe M, McLaughlin RE, Whiteaker JD, Roy PH, Alm RA, Corbeil J, Gardner H. 2015. The resistome of *Pseudomonas aeruginosa* in relationship to phenotypic susceptibility. Antimicrobial Agents and Chemotherapy 59:427.

34. Richardot C, Plesiat P, Fournier D, Monlezun L, Broutin I, Llanes C. 2015. Carbapenem resistance in cystic fibrosis strains of *Pseudomonas aeruginosa* as a result of amino acid substitutions in porin OprD. Int J Antimicrob Agents 45:529–32.

35. Chuanchuen R, Narasaki CT, Schweizer HP. 2002. The MexJK efflux pump of *Pseudomonas aeruginosa* requires OprM for antibiotic efflux but not for efflux of triclosan. J Bacteriol 184:5036–44.

36. Masuda N, Gotoh N, Ishii C, Sakagawa E, Ohya S, Nishino T. 1999. Interplay between chromosomal beta-lactamase and the MexAB-OprM efflux system in intrinsic resistance to beta-lactams in *Pseudomonas aeruginosa*. Antimicrob Agents Chemother 43:400–2.

37. Langaee TY, Gagnon L, Huletsky A. 2000. Inactivation of the *ampD* gene in *Pseudomonas aeruginosa* leads to moderate-basal-level and hyperinducible AmpC beta-lactamase expression. Antimicrob Agents Chemother 44:583–9.

38. Kong KF, Jayawardena SR, Indulkar SD, Del Puerto A, Koh CL, Hoiby N, Mathee K. 2005. *Pseudomonas aeruginosa* AmpR is a global transcriptional factor that regulates expression of AmpC and PoxB beta-lactamases, proteases, quorum sensing, and other virulence factors. Antimicrob Agents Chemother 49:4567–75.

39. Moya B, Dotsch A, Juan C, Blazquez J, Zamorano L, Haussler S, Oliver A. 2009. Betalactam resistance response triggered by inactivation of a nonessential penicillin-binding protein. PLoS Pathog 5:e1000353.

40. Tsutsumi Y, Tomita H, Tanimoto K. 2013. Identification of novel genes responsible for overexpression of *ampC* in *Pseudomonas aeruginosa* PAO1. Antimicrob Agents Chemother 57:5987–93.

41. Berrazeg M, Jeannot K, Ntsogo Enguene VY, Broutin I, Loeffert S, Fournier D, Plesiat P. 2015. Mutations in beta-lactamase AmpC increase resistance of *Pseudomonas aeruginosa* isolates to antipseudomonal cephalosporins. Antimicrob Agents Chemother 59:6248–55.

42. Han S, Zaniewski RP, Marr ES, Lacey BM, Tomaras AP, Evdokimov A, Miller JR, Shanmugasundaram V. 2010. Structural basis for effectiveness of siderophore-conjugated monocarbams against clinically relevant strains of *Pseudomonas aeruginosa*. Proc Natl Acad Sci U S A 107:22002–7.

43. Diaz Caballero J, Clark ST, Coburn B, Zhang Y, Wang PW, Donaldson SL, Tullis DE, Yau YCW, Waters VJ, Hwang DM, Guttman DS. 2015. Selective sweeps and parallel pathoadaptation drive *Pseudomonas aeruginosa* evolution in the cystic fibrosis lung. mBio 6:e00981–15.

44. Clark ST, Sinha U, Zhang Y, Wang PW, Donaldson SL, Coburn B, Waters VJ, Yau YCW, Tullis DE, Guttman DS, Hwang DM. 2019. Penicillin-binding protein 3 is a common adaptive target among *Pseudomonas aeruginosa* isolates from adult cystic fibrosis patients treated with beta-lactams. Int J Antimicrob Agents 53:620–628.

45. Jorth P, McLean K, Ratjen A, Secor PR, Bautista GE, Ravishankar S, Rezayat A, Garudathri J, Harrison JJ, Harwood RA, Penewit K, Waalkes A, Singh PK, Salipante SJ. 2017. Evolved aztreonam resistance is multifactorial and can produce hypervirulence in *Pseudomonas aeruginosa*. mBio 8:e00517–17.

46. Poole K, Tetro K, Zhao Q, Neshat S, Heinrichs DE, Bianco N. 1996. Expression of the multidrug resistance operon *mexA-mexB-oprM* in *Pseudomonas aeruginosa: mexR* encodes a regulator of operon expression. Antimicrob Agents Chemother 40:2021–8.

47. Choudhury D, Das Talukdar A, Dutta Choudhury M, Maurya AP, Paul D, Dhar Chanda D, Chakravorty A, Bhattacharjee A. 2015. Transcriptional analysis of MexAB-OprM efflux pumps system of *Pseudomonas aeruginosa* and its role in carbapenem resistance in a tertiary referral hospital in India. PLOS ONE 10:e0133842.

48. Cabot G, Zamorano L, Moya B, Juan C, Navas A, Blazquez J, Oliver A. 2016. Evolution of *Pseudomonas aeruginosa* antimicrobial resistance and fitness under low and high mutation rates. Antimicrob Agents Chemother 60:1767–78.

49. Castanheira M, Deshpande LM, Costello A, Davies TA, Jones RN. 2014. Epidemiology and carbapenem resistance mechanisms of carbapenem-non-susceptible *Pseudomonas L. aeruginosa* collected during 2009-11 in 14 European and Mediterranean countries. J Antimicrob Chemother 69:1804–14.

50. Jeannot K, Sobel ML, El Garch F, Poole K, Plesiat P. 2005. Induction of the MexXY efflux pump in *Pseudomonas aeruginosa* is dependent on drug-ribosome interaction. J Bacteriol 187:5341–6.

51. Prickett MH, Hauser AR, McColley SA, Cullina J, Potter E, Powers C, Jain M. 2017. Aminoglycoside resistance of *Pseudomonas aeruginosa* in cystic fibrosis results from convergent evolution in the *mexZ* gene. Thorax 72:40–47.

52. Guenard S, Muller C, Monlezun L, Benas P, Broutin I, Jeannot K, Plesiat P. 2014. Multiple mutations lead to MexXY-OprM-dependent aminoglycoside resistance in clinical strains of *Pseudomonas aeruginosa*. Antimicrob Agents Chemother 58:221–8.

53. Alguel Y, Lu D, Quade N, Sauter S, Zhang X. 2010. Crystal structure of MexZ, a key repressor responsible for antibiotic resistance in *Pseudomonas aeruginosa*. J Struct Biol 172:305–10.

54. Lau CH, Hughes D, Poole K. 2014. MexY-promoted aminoglycoside resistance in *Pseudomonas aeruginosa*: involvement of a putative proximal binding pocket in aminoglycoside recognition. MBio 5:e01068.

55. Vettoretti L, Plesiat P, Muller C, El Garch F, Phan G, Attree I, Ducruix A, Llanes C. 2009. Efflux unbalance in *Pseudomonas aeruginosa* isolates from cystic fibrosis patients. Antimicrob Agents Chemother 53:1987–97.

56. Lau CH-F, Fraud S, Jones M, Peterson SN, Poole K. 2013. Mutational activation of the AmgRS two-component system in aminoglycoside-resistant *Pseudomonas aeruginosa*. Antimicrobial agents and chemotherapy 57:2243–2251.

57. Park H, Inouye M. 1997. Mutational analysis of the linker region of EnvZ, an osmosensor in *Escherichia coli*. Journal of bacteriology 179:4382–4390.

58. Giraud A, Arous S, Paepe MD, Gaboriau-Routhiau V, Bambou J-C, Rakotobe S, Lindner AB, Taddei F, Cerf-Bensussan N. 2008. Dissecting the genetic components of adaptation of *Escherichia coli* to the mouse gut. PLOS Genetics 4:e2.

59. Feng Y, Jonker MJ, Moustakas I, Brul S, ter Kuile BH. 2016. Dynamics of Mutations during Development of Resistance by *Pseudomonas aeruginosa* against Five Antibiotics. Antimicrobial Agents and Chemotherapy 60:4229.

60. Bolard A, Plesiat P, Jeannot K. 2018. Mutations in gene *fusA1* as a novel mechanism of aminoglycoside resistance in clinical strains of *Pseudomonas aeruginosa*. Antimicrob Agents Chemother 62.

61. Lopez-Causape C, Cabot G, Del Barrio-Tofino E, Oliver A. 2018. The versatile mutational resistome of *Pseudomonas aeruginosa*. Front Microbiol 9:685.

62. Markussen T, Marvig RL, Gómez-Lozano M, Aanæs K, Burleigh AE, Høiby N, Johansen HK, Molin S, Jelsbak L. 2014. Environmental heterogeneity drives within-host diversification and evolution of *Pseudomonas aeruginosa*. mBio 5:e01592–14.

63. Greipel L, Fischer S, Klockgether J, Dorda M, Mielke S, Wiehlmann L, Cramer N, Tümmler B. 2016. Molecular epidemiology of mutations in antimicrobial resistance loci of *Pseudomonas aeruginosa* isolates from airways of cystic fibrosis patients. Antimicrobial Agents and Chemotherapy 60:6726.

64. Chung JCS, Becq J, Fraser L, Schulz-Trieglaff O, Bond NJ, Foweraker J, Bruce KD, Smith GP, Welch M. 2012. Genomic variation among contemporary *Pseudomonas aeruginosa L.* isolates from chronically infected cystic fibrosis patients. Journal of Bacteriology 194:4857.

65. Chavez-Jacobo VM, Hernandez-Ramirez KC, Romo-Rodriguez P, Perez-Gallardo RV, Campos-Garcia J, Gutierrez-Corona JF, Garcia-Merinos JP, Meza-Carmen V, Silva-Sanchez J, Ramirez-Diaz MI. 2018. CrpP is a novel ciprofloxacin-modifying enzyme encoded by the *Pseudomonas aeruginosa* pUM505 plasmid. Antimicrob Agents Chemother 62.

66. Lopez-Causape C, Oliver A. 2017. Insights into the evolution of the mutational resistome of *Pseudomonas aeruginosa* in cystic fibrosis. Future Microbiol 12:1445–1448.

67. Mulet X, Macia MD, Mena A, Juan C, Perez JL, Oliver A. 2009. Azithromycin in *Pseudomonas aeruginosa* biofilms: bactericidal activity and selection of *nfxB* mutants. Antimicrob Agents Chemother 53:1552–60.

68. Miller AK, Brannon MK, Stevens L, Johansen HK, Selgrade SE, Miller SI, Hoiby N, Moskowitz SM. 2011. PhoQ mutations promote lipid A modification and polymyxin resistance of *Pseudomonas aeruginosa* found in colistin-treated cystic fibrosis patients. Antimicrob Agents Chemother 55:5761–9.

69. Lee JY, Ko KS. 2014. Mutations and expression of PmrAB and PhoPQ related with colistin resistance in *Pseudomonas aeruginosa* clinical isolates. Diagn Microbiol Infect Dis 78:271–6.

70. Fernandez L, Jenssen H, Bains M, Wiegand I, Gooderham WJ, Hancock RE. 2012. The two-component system CprRS senses cationic peptides and triggers adaptive resistance in *Pseudomonas aeruginosa* independently of ParRS. Antimicrob Agents Chemother 56:6212–22.

71. Gutu AD, Sgambati N, Strasbourger P, Brannon MK, Jacobs MA, Haugen E, Kaul RK, Johansen HK, Hoiby N, Moskowitz SM. 2013. Polymyxin resistance of *Pseudomonas aeruginosa phoQ* mutants is dependent on additional two-component regulatory systems. Antimicrob Agents Chemother 57:2204–15.

72. Johansen HK, Moskowitz SM, Ciofu O, Pressler T, Hoiby N. 2008. Spread of colistin resistant non-mucoid *Pseudomonas aeruginosa* among chronically infected Danish cystic fibrosis patients. J Cyst Fibros 7:391–7.

73. Lo Sciuto A, Imperi F. 2018. Aminoarabinosylation of Lipid A is critical for the development of colistin resistance in *Pseudomonas aeruginosa*. Antimicrob Agents Chemother 62.

74. Muller C, Plesiat P, Jeannot K. 2011. A two-component regulatory system interconnects resistance to polymyxins, aminoglycosides, fluoroquinolones, and beta-lactams in *Pseudomonas aeruginosa*. Antimicrob Agents Chemother 55:1211–21.

75. Marvig RL, Søndergaard MSR, Damkiær S, Høiby N, Johansen HK, Molin S, Jelsbak L. 2012. Mutations in 23S rRNA confer resistance against azithromycin in *Pseudomonas aeruginosa*. Antimicrobial agents and chemotherapy 56:4519–4521.

76. Eliopoulos GM, Blázquez J. 2003. Hypermutation as a Factor Contributing to the Acquisition of Antimicrobial Resistance. Clinical Infectious Diseases 37:1201–1209.

77. Taddei F, Radman M, Maynard-Smith J, Toupance B, Gouyon PH, Godelle B. 1997. Role of mutator alleles in adaptive evolution. Nature 387:700–2.

78. Tenaillon O, Toupance B, Le Nagard H, Taddei F, Godelle B. 1999. Mutators, population size, adaptive landscape and the adaptation of asexual populations of bacteria. Genetics 152:485–93.

79. Sniegowski PD, Gerrish PJ, Johnson T, Shaver A. 2000. The evolution of mutation rates: separating causes from consequences. Bioessays 22:1057–66.

80. Andre JB, Godelle B. 2006. The evolution of mutation rate in finite asexual populations. Genetics 172:611–26.

81. MacLean RC, Perron GG, Gardner A. 2010. Diminishing returns from beneficial mutations and pervasive epistasis shape the fitness landscape for rifampicin resistance in *Pseudomonas aeruginosa*. Genetics 186:1345–54.

82. MacLean RC, Vogwill T. 2014. Limits to compensatory adaptation and the persistence of antibiotic resistance in pathogenic bacteria. Evol Med Public Health 2015:4–12.

83. Wong A. 2017. Epistasis and the Evolution of Antimicrobial Resistance. Frontiers in Microbiology 8.

84. Lind PA, Libby E, Herzog J, Rainey PB. 2019. Predicting mutational routes to new adaptive phenotypes. eLife 8:e38822.

85. Huse HK, Kwon T, Zlosnik JEA, Speert DP, Marcotte EM, Whiteley M. 2010. Parallel Evolution in *Pseudomonas aeruginosa* over 39,000 Generations *In Vivo*. mBio 1:e00199–10.

86. Wong A, Kassen R. 2011. Parallel evolution and local differentiation in quinolone resistance in *Pseudomonas aeruginosa*. Microbiology 157:937–44.

87. Zhanel GG, Chung P, Adam H, Zelenitsky S, Denisuik A, Schweizer F, Lagace-Wiens PR, Rubinstein E, Gin AS, Walkty A, Hoban DJ, Lynch JP, 3rd, Karlowsky JA. 2014. Ceftolozane/tazobactam: a novel cephalosporin/beta-lactamase inhibitor combination with activity against multidrug-resistant gram-negative bacilli. Drugs 74:31–51.

88. Cabot G, Bruchmann S, Mulet X, Zamorano L, Moya B, Juan C, Haussler S, Oliver A. 2014. *Pseudomonas aeruginosa* ceftolozane-tazobactam resistance development requires multiple mutations leading to overexpression and structural modification of AmpC. Antimicrob Agents Chemother 58:3091–9.

89. Desai MM, Fisher DS, Murray AW. 2007. The Speed of Evolution and Maintenance of Variation in Asexual Populations. Current Biology 17:385–394.

90. Sniegowski PD, Gerrish PJ. 2010. Beneficial mutations and the dynamics of adaptation in asexual populations. Philosophical Transactions of the Royal Society B: Biological Sciences 365:1255–1263.

91. Kumar P, Henikoff S, Ng PC. 2009. Predicting the effects of coding non-synonymous variants on protein function using the SIFT algorithm. Nat Protoc 4:1073–81.

92. Choi Y, Sims GE, Murphy S, Miller JR, Chan AP. 2012. Predicting the functional effect of amino acid substitutions and indels. PLoS One 7:e46688.

93. Hecht M, Bromberg Y, Rost B. 2015. Better prediction of functional effects for sequence variants. BMC Genomics 16:S1.

94. Zankari E, Hasman H, Cosentino S, Vestergaard M, Rasmussen S, Lund O, Aarestrup FM, Larsen MV. 2012. Identification of acquired antimicrobial resistance genes. J Antimicrob Chemother 67:2640–4.

95. CLSI. 2019. Performance Standards for Antimicrobial Susceptibility Testing. 29th ed. Wayne, PA: Clinical and Laboratory Standards Institute.

96. Barnes MD, Taracila MA, Rutter JD, Bethel CR, Galdadas I, Hujer AM, Caselli E, Prati F, Dekker JP, Papp-Wallace KM, Haider S, Bonomo RA. 2018. Deciphering the evolution of cephalosporin resistance to ceftolozane-tazobactam in *Pseudomonas aeruginosa*. mBio 9:e02085–18.

