## Supplemental Figures and Tables for "Hypermutator *Pseudomonas aeruginosa* exploits multiple genetic pathways to develop multidrug resistance during long-term infections in the airways of cystic fibrosis patients"

**SUPPLEMENTARY MATERIAL**

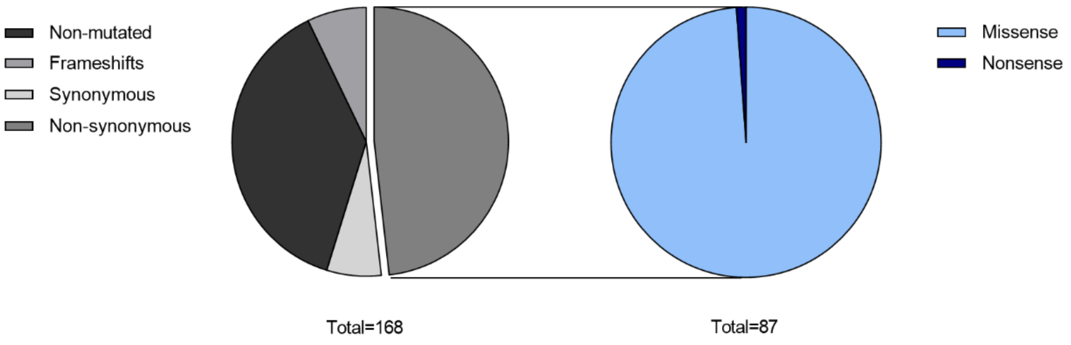


**Figure S1. Analysis of type of mutations found in antibiotic resistance genes in *P. aeruginosa* isolates from CFD patient.**

Pie charts indicate the observed percentage for each kind of mutation respect to the total number of mutations occurring in the 168 belonging to the *P. aeruginosa* resistome.

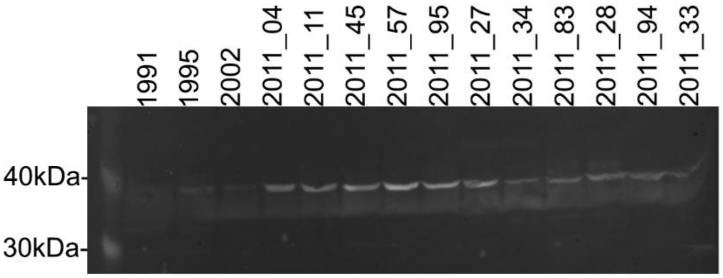


**Figure S2. Western blot of CFD isolates.**

Total proteins (25µg) were obtained from whole-cell lysates from each *P. aeruginosa* clinical isolates, resolved in a 12% polyacrylamide gel, and tested with a PDC-3 antibody.

**Table S1. Nonsynonymous and frameshift mutations found within 93 out of 168 antibiotic resistance genes in CFD collection.**

| Gene locus | Gene name | Mutation (no. Of isolates) |
|---|---|---|
| PA0004 | *gyrB* | S618L (2) |
| PA0005 | *lptA* | I7T (1) |
| PA0301 | *spuE* | A335V (11) |
| PA0302 | *spuF* | A216V (1), D122G (1), H321R (5) |
| PA0425 | *mexA* | A6V (1) |
| PA0426 | *mexB* | A562V (1), M626V (5) |
| PA0427 | *oprM* | T477A (1), -1C at 1447 (1) |
| PA0464 | *creC* | D73G (1) |
| PA0486 | *yihE* | Y39H (5), R136Q (1) |
| PA0779 | *asrA* | A375T (5) |
| PA0807 | *ampDh3* | Q65R (2), K129R (1) |
| PA0958 | *oprD* | S278P (1) |
| PA1179 | *phoP* | V30A (11) |
| PA1345 | *gshB* | V100M (1) |
| PA1409 | *aphA* | N174S (11) |
| PA1430 | *lasR* | Y69C (1) |
| PA1588 | *sucC* | E249G (11) |
| PA1797 | - | Y77H (1) |
| PA1799 | *parR* | A45T (1) |
| PA1801 | *clpP* | L208P (10) |
| PA1803 | *lon* | A644T (5) |
| PA1812 | *mltD* | L527P (11) |
| PA1886 | *polB* | T161I (1), A374T (5), G509S (12) |
| PA2006 | - | P130T (1) |
| PA2018 | *mexY* | S476P (1), G815S (11), G855S (2), I919T (1), V979A (5), E1043K (1) |
| PA2019 | *mexX* | A115V (11), T144I (1), +1G at 32 (1) |
| PA2020 | *mexZ* | V29A (1) |
| PA2071 | *fusA2* | V480I (1), A598T (1), -1C at 268 (5) |
| PA2272 | PBP3a | V234A (2), I254T (1) |
| PA2490 | *ydbB* | V45A (5) |
| PA2492 | *mexT* | F34L (10) |
| PA2494 | *mexF* | A54T (5), V133I (5), L182F (1), T883I (1), -CAT at 1308/10 (1) |
| PA2495 | *oprN* | A195T (10), R210Q (5), A226T (2) |
| PA2522 | *czcC* | R293L (1) |
| PA2523 | *czcR* | V48A (10), D180N (1) |

| | | |
|---|---|---|
| PA2525 | *opmB* | L146F (1) |
| PA2526 | *muxC* | Y869C (1) |
| PA2527 | *muxB* | Y817C (5), P943L (1) |
| PA2528 | *muxA* | T231A (1), T358A (10) |
| PA2642 | *nuoG* | A480T (10) |
| PA2798 | - | A177D (5) |
| PA3005 | *nagZ* | R303C (1) |
| PA3013 | *foaB* | E186G (1) |
| PA3050 | *pyrD* | R89Q (1) |
| PA3078 | *cprS* | V226A (2), G396S (10) |
| PA3141 | *capD* | L280P (13) |
| PA3168 | *gyrA* | T83I (1), D87G (13) |
| PA3522 | *mexQ* | -1C at 1516 (5) |
| PA3523 | *mexP* | +1C at 1069 (1) |
| PA3602 | *yerD* | Y120H (2), D206G (10) |
| PA3676 | *mexK* | V216A (5), H498Y (2), D857G (1) |
| PA3721 | *nalC* | M151T (2) |
| PA4020 | *mpl* | +1G at 1244 (13) |
| PA4069 | - | A232V (2) |
| PA4110* | *ampC* | A89V (10), Q120K (10), P154L (1), H189Y (2), V213A (11), G216S (1), N321S (8), V330I (1) |
| PA4207 | *mexI* | Y140C (2), E461K (5), E339K (1) |
| PA4208 | *opmD* | A75V (1) |
| PA4218 | *ampP* | -1G at 649 (4) |
| PA4260 | *rplB* | G138A (13) |
| PA4266 | *fusA1* | V93A (1), V338A (1), A481V (2), A595V (5), D588G (1), Y683C (1) |
| PA4269 | *rpoC* | T1093A (1), T1328A (1) |
| PA4270 | *rpoB* | E477G (2) |
| PA4280.2 | - | A→G at 2044 (1) |
| PA4315 | *mvaT* | -GC at 229 (1) |
| PA4374 | *mexV* | Y123H (1) |
| PA4380 | *colS* | T138A (2) |
| PA4381 | *colR* | D138N (11) |
| PA4393 | *ampG* | L114P (10), R371H (1) |
| PA4406 | *lpxC* | Y230H (1) |
| PA4418 | PBP3 | G63D (10), P143S (1), Y367C (5), H394R (5), N427S (1), Q458R (2), G469S (1), Q475R (1), R504L (5), P512L (1), V523M (2), V523A (5), F533L (11) |
| PA4444 | *mltB1* | G308D (5) |
| PA4521 | *ampE* | A170T (5) |
| PA4522 | *ampD* | E186G (2) |
| PA4597 | *oprJ* | S198N (1) |
| PA4598 | *mexD* | +1C at 1738 (1), N213S (2), W1023STOP (5) |
| PA4599 | *mexC* | P365S (3) |

| | | |
|---|---|---|
| PA4661 | *pagL* | G67D (1), R146W (1) |
| PA4671 | *rplY* | Q41R (1) |
| PA4690.2 | - | C→T at 2597 (10) |
| PA4700 | PBP1b | A107T (10), I245V (1) |
| PA4751 | *ftsH* | R181C (2) |
| PA4774 | - | V132M (1) |
| PA4777 | *pmrB* | T132A (10), V431I (1) |
| PA4964 | *parC* | P308L (1), T705A (1) |
| PA4967 | *parE* | V199M (12), D462G (10), S492F (1) |
| PA5045 | PBP1a | G411D (1) |
| PA5117 | *typA* | P514L (1) |
| PA5199 | *amgS* | H88R (5), P116L (1), A203V (10), T394A (1) |
| PA5235 | *glpT* | L433P (12) |
| PA5297 | *poxB* | V33A (2), A161T (2), R489S (1) |
| PA5332 | *crc* | V102A (10) |
| PA5528 | - | Q97R (1) |
| PA5542 | - | -1C at 738 (1), H277R (2) |

*Pseudomonas aeruginosa* PAO1 was used as reference genome for annotation of the 168 antibiotic resistance-related genes.

*Numbering of amino acids refers to the mature protein of PAO1 strain, after cleavage of 26 N-terminal amino acid residues from the signal peptide.

**Table S2. Number of genes mutated and type of mutations found in the sequenced genomes.**

| Genome | Genes[a] | Missense | Nonsense | Synonymous | dN/dS ratio[b] |
|---|---|---|---|---|---|
| 1995 | 22 | 25 | 0 | 6 | 1.39 |
| 2002 | 25 | 26 | 0 | 6 | 1.44 |
| 2011_95 | 47 | 62 | 1 | 19 | 1.11 |
| 2011_11 | 49 | 62 | 1 | 19 | 1.11 |
| 2011_04 | 47 | 67 | 1 | 19 | 1.19 |
| 2011_45 | 49 | 65 | 1 | 20 | 1.16 |
| 2011_57 | 46 | 64 | 1 | 19 | 1.14 |
| 2011_27 | 42 | 52 | 0 | 13 | 1.33 |
| 2011_34 | 43 | 51 | 0 | 17 | 1 |
| 2011_83 | 36 | 43 | 0 | 20 | 0.72 |
| 2011_28 | 43 | 58 | 0 | 18 | 1.07 |
| 2011_94 | 42 | 57 | 0 | 14 | 1.36 |
| 2011_33 | 36 | 41 | 0 | 14 | 0.98 |

Mutations were scored using CFD_1991 as the reference genome. [a]To calculate mutated genes, each gene was counted as 1 if they harbored one or more mutations. [b]To calculate selection coefficients (dN/dS ratio), we assumed identical codon usages to strain PAO1, in which 25% of random mutations are synonymous (L. Yang, L. Jelsbak, R. L. Marvig, S. Damkiaer, C. T. Workman, M. H. Rau, S. K. Hansen, A. Folkesson, H. K. Johansen, O. Ciofu, N. Hoiby, M. O. Sommer and S. Molin, PNAS 108 (18) 7481-7486, 2011, https://doi:10.1073/pnas.1018249108).
